# Epigenome-wide association meta-analysis of DNA methylation with coffee and tea consumption

**DOI:** 10.1101/2020.04.15.042267

**Authors:** Irma Karabegović, Eliana Portilla-Fernandez, Yang Li, Jiantao Ma, Silvana C.E. Maas, Daokun Sun, Emily A. Hu, Brigitte Kühnel, Yan Zhang, Srikant Ambatipudi, Giovanni Fiorito, Jian Huang, Juan E. Castillo-Fernandez, Kerri L. Wiggins, Niek de Klein, Sara Grioni, Brenton R. Swenson, Silvia Polidoro, Jorien L. Treur, Cyrille Cuenin, Pei-Chien Tsai, Ricardo Costeira, Veronique Chajes, Kim Braun, Niek Verweij, Anja Kretschmer, Lude Franke, Joyce B.J. van Meurs, André G. Uitterlinden, Robert J. de Knegt, M. Arfan Ikram, Abbas Dehghan, Annette Peters, Ben Schöttker, Sina A. Gharib, Nona Sotoodehnia, Jordana T. Bell, Paul Elliott, Paolo Vineis, Caroline Relton, Zdenko Herceg, Hermann Brenner, Melanie Waldenberger, Casey M. Rebholz, Trudy Voortman, Qiuwei Pan, Myriam Fornage, Daniel Levy, Manfred Kayser, Mohsen Ghanbari

## Abstract

Coffee and tea are extensively consumed beverages worldwide. Observational studies have shown contradictory findings for the association between consumption of these beverages and different health outcomes. Epigenetics is suggested as a mechanism mediating the effects of dietary and lifestyle factors on disease onset. We conducted epigenome-wide association studies (EWAS) on coffee and tea consumptions in 15,789 participants of European and African-American ancestries from 15 cohorts. EWAS meta-analysis revealed 11 CpG sites significantly associated with coffee consumption (*P*-value <1.1×10^-7^), nine of them annotated to the genes *AHRR, F2RL3, FLJ43663, HDAC4, GFI1* and *PHGDH*, and two CpGs suggestively associated with tea consumption (*P*-value<5.0×10^-6^). Among these, cg14476101 was significantly associated with expression of its annotated gene *PHGDH* and risk of fatty liver disease. Knockdown of *PHGDH* expression in liver cells showed a correlation with expression levels of lipid-associated genes, suggesting a role of *PHGDH* in hepatic-lipid metabolism. Collectively, this study indicates that coffee consumption is associated with differential DNA methylation levels at multiple CpGs, and that coffee-associated epigenetic variations may explain the mechanism of action of coffee consumption in conferring disease risk.

## Introduction

Excluding water, coffee and tea are the most commonly consumed beverages around the world and preference for one or the other varies between individuals as well as in populations^1^. Both coffee and tea are sources of complex compounds with different chemical classes, the most commonly known is caffeine^2^. Caffeine belongs to the methylxanthines family, which consists of frequently ingested pharmacologically active substances, for example, through the stimulation of the central nervous system^3^. Although caffeine is present in both coffee and tea, its concentration in tea is much lower^4^. Moreover, both beverages differ on the bioavailability of polyphenols and other chemical compounds^5^. The biochemistry of coffee and tea has been extensively documented, indicating that different roasting, temperatures, or brewing of the two can impact the abundancy and bioavailability of their complex compounds^6,7^. There has been an ongoing debate as to whether habitual consumption of coffee^8^ and tea^9^ is beneficial or harmful to health. The conclusion varies among outcomes; lowering the risk for type 2 diabetes and cardiovascular diseases^10,11^, and increasing serum levels of low-density lipoprotein (LDL) and total cholesterol^12^. Given the presence of different compounds in these two beverages with diverse effects on disease risk, observational studies have given rise to contradictory findings, especially regarding coffee consumption^8^. Previous studies, however, have shown a consistent association of coffee consumption with lower risk of overall mortality^13^ and liver diseases ^14,15^. Yet, the biological mechanisms underlying associations of coffee and tea consumptions with disease risk remain to be investigated.

Epigenetics represents modifications to DNA that do not change the underlying DNA sequence, but instead, influence gene expression^16^. The most extensively studied epigenetic mechanism is DNA methylation, where a methyl group (-CH3) is added or removed to the cytosine nucleotide that is followed by a guanine nucleotide in the DNA sequence, known as Cytosine-Phosphate Guanine (CpG)^16^, resulting in altered gene expression. DNA methylation levels differ by age, sex and lifestyle factors, including dietary exposures^17–19^. Here, we postulate that alteration of DNA methylation via coffee or tea consumptions is an underlying mechanism linking the intake of these beverages to health outcomes. Previous epigenome-wide association studies (EWAS) have reported suggestive association of a few CpGs with tea or coffee consumptions^20,21^; however, these studies were limited by modest sample sizes. In the present study, we conducted large-scale EWAS meta-analyses on coffee and tea consumptions in 15,789 participants of European and African-American ancestries from 15 cohort studies. The associated CpG were analyzed to evaluate their correlations with genetic variation and gene expression. Additionally, we explored the potential causal effect of coffee consumption on the identified CpGs and different health outcomes and performed experimental studies for a candidate gene to investigate its link to liver diseases.

## Methods

### Study population

**Figure 1** depicts an overview of the study flow. This study was conducted within the framework of the Cohort for Heart and Aging Research in Genomic Epidemiology (CHARGE consortium)^22^ and additional participating cohorts, resulting in a total sample size of 15,789 participants. Clinical characteristics of the participants included in our study are presented in **Table 1**. The discovery phase included 9,612 participants of European ancestry (EA) from the following cohorts (listed in alphabetical order): Airwave^23^, Avon Longitudinal Study of Parents and Children (ALSPAC)^24^, two independent datasets from the ESTHER Study (ESTHER_a and ESTHER_b)^25^, Framingham Heart Study (FHS)^26^, Cooperative Health Research in the Augsburg Region Study (KORA)^27^, two cohorts of the Rotterdam Study (RS-II and RS-III)^28^ and TwinsUK^29^. We sought replication of the associated CpG sites from the discovery phase, in an independent population consisting of 6,177 participants of European and African American (AA) ancestries (18.3%). The replication phase included two ethnically different sub cohorts of Atherosclerosis Risk in Communities Study (ARIC_EA and ARIC_AA)^30^, two ethnically different sub cohorts from the Cardiovascular Health Study (CHS_EA and CHS_AA)^31^, and two independent studies from the European Prospective Investigation into Cancer and Nutrition (EPIC_Italy and EPIC_IARC)^32^. All participants provided written informed consent, and all contributing cohorts confirmed compliance with their local research ethics committees or Institutional Review Boards. Detailed information of the participating cohorts are provided in Supplementary Information.

**Figure 1.**
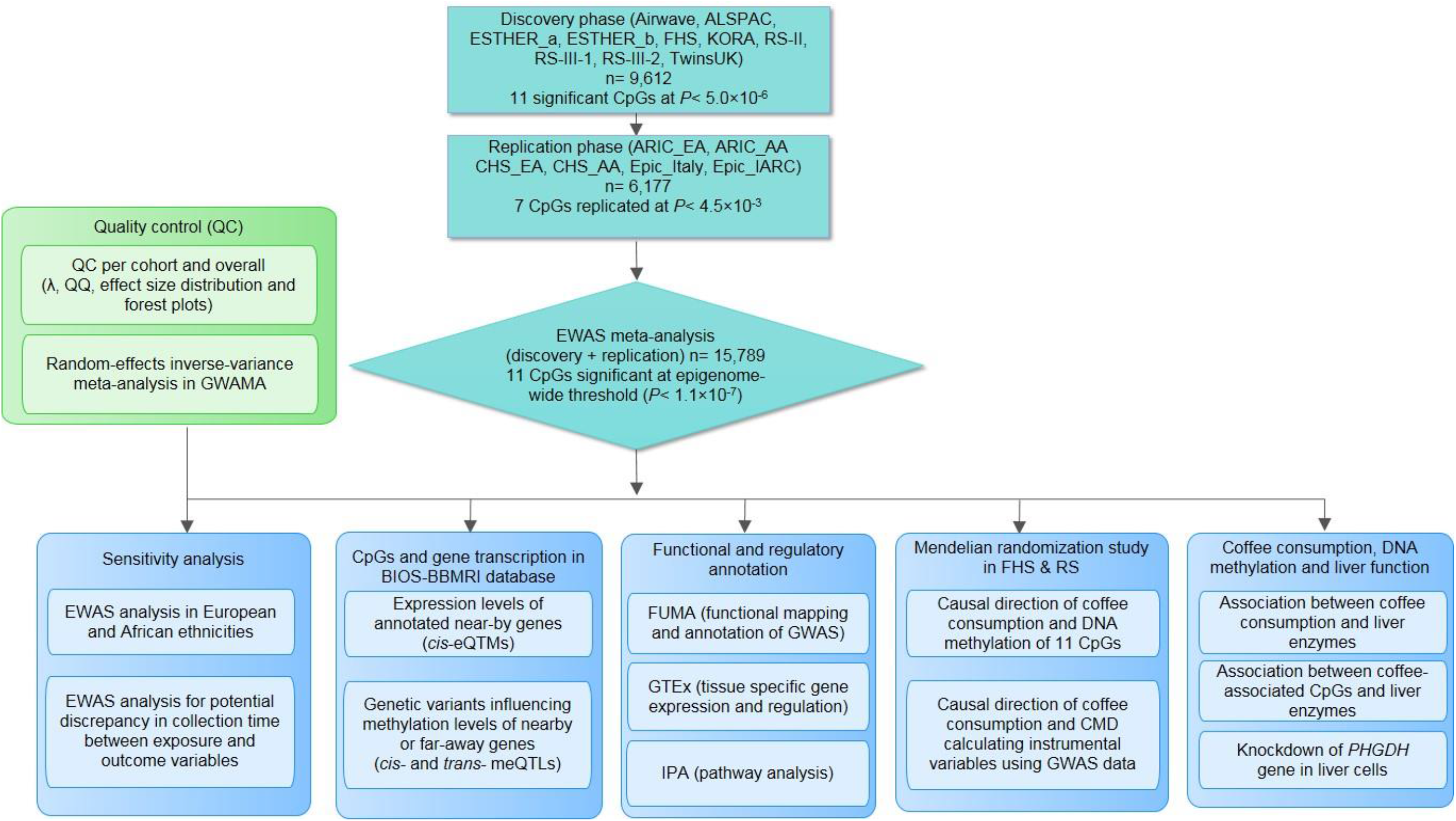
An overview of our study including EWAS meta-analysis to identify DNA methylation sites associated with coffee and tea consumption, and post-EWAS *in silico* and in vitro experiments. RS= Rotterdam Study, FHS= Framingham Heart Study, ALSPAC= The Avon Longitudinal Study of Parents and Children, CHS= Cardiovascular Health Study, ARIC= The Atherosclerosis Risk in Communities, EPIC= Prospective Investigation into Cancer and Nutrition, KORA= Cooperative Health Research in the Augsburg Region Study.

**Table 1:** Characteristics of the cohort participants

| Cohort | N | Ethnicity | Age (years) | Women (%) | Coffee (cups/day) | Tea (cups/day) | Smoking |  |  | BMI (kg/m<sup>2</sup>) | Alcohol (gr/day) |
| --- | --- | --- | --- | --- | --- | --- | --- | --- | --- | --- | --- |
|  |  |  |  |  |  |  | current | former | never |  |  |
| **Discovery phase** |  |  |  |  |  |  |  |  |  |  |  |
| RS-III-2 | 149 | EA | 61.01 (4.42) | 116 (66.3%) | 3.52 (2.22) | 1.43 (1.5) | 20 (11.4%) | 89 (50.9%) | 66 (37.7%) | 27.47 (4.13) | 8.18 (7) |
| RS-II-3 | 367 | EA | 70.97 (3.21) | 242 (55.09%) | 2.85 (1.78) | 1.18 (1.3) | 38 (8.8%) | 243 (56.1%) | 152 (35.1%) | 27.69 (4.10) | 8.55 (7.55) |
| RS-III-1 | 549 | EA | 58.89 (7.8) | 283 (51.5%) | 3.49 (2.13) | 1.23 (1.31) | 148 (27%) | 235 (42.8%) | 166 (30.2%) | 27.27 (4.65) | 9.14 (8.68) |
| ALSPAC | 331 | EA | 48.04 (4.03) | 331 (100%) | 1.69 (1.65) | 3.02 (2.08) | current + former = 152 (46%) |  | 179 (54%) | 25.67 (4.67) | 6.5 (6.66) |
| KORA | 1535 | EA | 54 (8.8) | 774 (50.4%) | 3.4 (3) | 1.1 (2.1) | 307 (20%) | 568 (37%) | 660 (43%) | 27.7 (4.48) | 17.3 (22.87) |
| FHS | 3718 | EA | 59 (13) | 2008 (54%) | 1.35 (1.28) | 0.37 (0.77) | 297 (8%) | never + former = 3421 (92%) |  | 28 (5.40) | 10.80 (15.50) |
| ESTHER_A | 973 | EA | 62 (6.5) | 488 (50.1%) | 1.7 (1.1) | 0.4 (0.7) | 181 (18.9%) | 315 (32.8%) | 464 (48.3%) | 27.8 (4.2) | 12.3 (15.4) |
| ESTHER_B | 532 | EA | 62 (6.6) | 327 (61.5%) | 1.7 (1.1) | 0.5 (0.8) | 96 (18.5%) | 177 (34.1%) | 426 (47.4%) | 27.6 (4.40) | 10.9 (14.50) |
| TwinsUK | 552 | EA | 58.45 (9.97) | 552 (100%) | 2.37 (2.46) | 3.35 (2.94) | 66 (12%) | 149 (27%) | 337 (61%) | 26.51 (4.94) | 6.37 (9.15) |
| Airwave | 906 | EA | 41.6 (9.3) | 380 (41.9%) | 1.8 (2) | 3.1 (2.6) | 93 (10.3%) | 217 (24%) | 596 (65.8%) | 27.2 (4.50) | (never, current, ex) |
| Lifelines | 186 | EA | 45.04 (13.6) | 160 (57%) | 3.94 (2.03) | N/A | 34 (12.1%) | 99 (35.2%) | 148 (59.9%) | 25.78 (4.30) | 8.52 (9.42) |
| **Replication phase** |  |  |  |  |  |  |  |  |  |  |  |
| ARIC_EA | 1099 | EA | 57.73 (6.02) | 641 (58.33%) | 1.94 (1.99) | 0.77 (1.14) | 232 (21.11%) | 379 (34.49%) | 488 (44.40%) | 26.08 (4.40) | 5.34 (12.07) |
| ARIC_AA | 2736 | AA | 54.11 (6.12) | 1748 (63.89%) | 0.99 (1.33) | 0.38 (0.71) | 760 (27.78%) | 681 (24.89%) | 1295 (47.33%) | 29.89 (6.14) | 4.49 (14.07) |
| CHS_EA | 195 | EA | 78.6 (4.67) | 120 (61.5%) | 0.97 (1.33) | 0.57 (0.97) | 92 (47.2%) | 13 (6.6%) | 85 (43.6%) | 26.8 (4.61) | 2.27 (7.69) |
| CHA_AA | 185 | AA | 75.9 (4.71) | 121 (65.4%) | 0.6 (0.97) | 0.35 (0.62) | 79 (42.7%) | 19 (10.2%) | 80 (43.2%) | 28.3 (4.99) | 1.27 (4.23) |
| EPIC_Italy | 1096 | EA | 53.61 (6.95) | 761 (69%) | 2.79 (1.74) | 0.29 (0.61) | 299 (27%) | 296 (27%) | 501 (46%) | 25.98 (4.07) | 14.5 (18.79) |
| EPIC_IARC | 866 | EA | 52.23 (8.97) | 866 (100%) | 1.25 (1.18) | 0.69 (1.13) | 189 (22%) | 186 (21%) | 491 (57%) | 25.8 (4.45) | 9.1 (12.5) |

RS=Rotterdam Study. ALSPAC=The Avon Longitudinal Study of Parents and Children. KORA= Cooperative Health Research in the Augsburg Region Study. FHS = Framingham Heart Study.
ARIC= The Atherosclerosis Risk in Communities. CHS = Cardiovascular Health Study. EPIC = Prospective Investigation into Cancer and Nutrition. EA=European ancestry. AA=African ancestry

### Assessment of coffee and tea consumption

Data on coffee and tea intake was collected either by interview or food frequency questionnaires (FFQs). As some FFQs collected beverage intake over different periods of time (monthly, weekly or daily), data was harmonized among the cohorts to cups per day by taking the average intake of coffee/tea over the period of time specified by the FFQ utilized by each cohort. For instance, if the consumption was collected over the period of one month, daily consumption was estimated from the available data and multiplied with the frequency of consumption. Furthermore, if the beverage intake data was collected categorically, the median value was taken from the available data (e.g. 2.5 cups/day was used for the 2-3 cups/day category). If applicable, we excluded herbal tea and others, as green and black tea are derived from the different processing and harvesting of leaves from the same plant - *Camellia sinensis^9^.* Herbal tea does not contain any caffeine and green tea contains approximately half the caffeine compared to black tea (3.1 mg/fluid ounce, 5.9 mg/fluid ounce)^33^. In a subset of cohorts (RS-III-2, ALSPAC, EPIC_IARC, CHS_EA and CHS_AA), coffee and tea consumption data were collected a few years prior to the collection of whole blood, from which DNA methylation data was measured. Due to evidence from a previous research showing that coffee and tea consumptions tend to be stable over longer periods of time^34^, we used these data for our analysis.

### DNA methylation profiling

All participating cohorts measured DNA methylation in peripheral blood using the Infinium Human Methylation 450K Bead-Chip (Ilumina, San Diego, CA, USA) except Airwave cohort, where the Infinium Methylation EPIC (850K) Bead-Chip was used^35^. DNA methylation status was calculated with the β-value, signal from the methylated probe divided by the overall signal intensity. The methylation percentage of CpG sites was reported as a continuous β-value range between 0 (no methylation) and 1 (full methylation). Additional details are outlined in Supplementary Information. Cohort specific methods of normalization are shown in **Table S1**.

### EWAS of coffee and tea consumption

DNA methylation was considered as the dependent variable with coffee or tea consumption each as predictors of interest. Conventionally, each participating cohort performed an EWAS as a set of mixed effects linear-regression models, one CpG site at a time. In total, two linear mixed effects regression models were computed for each of the two exposures of interest. In the basic model (Model 1): we included age, sex, smoking status (never, former, and current), white blood cells (either measured or imputed based on the Houseman algorithm^36^ as fixed effects and technical covariates as random effects to control for batch effects. In the second model (Model 2), we additionally adjusted for body mass index (BMI, kg/m^2^) and alcohol consumption (g/day). The findings from Model 2 were considered as the primary results, as it is the most conservative model. All potential confounders were collected at the same time point of blood sampling for DNA methylation. Genetic principal components were included as covariates to account for population stratification if required. A detailed description of the covariates included in the models by each cohort is provided in Supplementary Information.

### EWAS meta-analysis

Since data originated from multiple sources, we performed quality control (QC) centrally. Each participating cohort submitted the EWAS summary statistics for the QC followed by meta-analysis. For this step, we used a specific package developed for QC within EWAS, namely “QCEWAS”^37^. We computed the genomic inflation factor (lambda) and checked quantile-quantile (QQ) plots for Model 2 for both coffee and tea consumption EWASs. Additionally, we computed effect-size distribution plots to assess the effect-size scale of each participating cohort. Results across independent cohorts were combined in both discovery and replication phase by using inverse variance fixed effects metaanalysis implemented in METAL v.2011-03-25^38^. Moreover, we assessed heterogeneity of effect estimates among cohorts using Cochran’s Q-test for heterogeneity implemented in METAL^38^. If there was nominal evidence for heterogeneity (*P* < 0.05), we performed random-effects inverse-variance meta-analysis using the method implemented in GWAMA^39^.

The discovery EWAS meta-analysis was conducted in 9,612 EA participants and differentially methylated CpGs at the suggestive threshold of *P* < 5.0×10^-6^ were interrogated. The CpGs that passed this threshold were tested for replication in the independent panel comprising 6,177 participants of European and African American ancestries, using the same models as implemented in the discovery phase and with a Bonferroni corrected p-value threshold, defined as 0.05 divided by the number of associated CpGs in the discovery phase. The significantly associated CpGs were retrieved from the combined meta-analysis with the whole samples at the epigenome-wide significance threshold (*P* < 1.0×10^-7^). If the CpG was missing from more than 4 participating cohorts, it was removed. Forest plots of the study-specific effect estimates were computed for significantly associated CpG sites.

Due to potential discrepancy in DNA methylation patterns between different ethnicities, we conducted meta-analysis EWASs separately in EA (n=12,868) and AA (n=2,921) participants. We first examined whether the significantly associated methylation sites in EA participants pass the Bonferroni-corrected p-value threshold in AA participants. Next, we tested whether the significant methylation sites in AA participants replicated in the meta-analysis of the EA participants.

Furthermore, we examined the potential impact of time varying exposure in cohorts that had different time points for methylation and coffee/tea consumption data collection. For this analysis, we excluded four cohorts (RS-III-2, ALSPAC, CHS_EA and CHS_AA) with different time points of data collection from the overall sample and meta-analyzed the remaining cohorts.

### Integration of EWAS results with genetic variation and gene expression

DNA methylation may have an impact on the transcription of genes, hence we used genetic variants and gene expression data from five Dutch biobanks (BIOS-BBMRI database) in a total of 3,841 whole blood samples (http://www.genenetwork.nl/biosqtlbrowser/), and explored whether DNA methylation levels of the significant CpGs affect expression levels of their annotated/nearby genes (*cis*-expression quantitative trait methylation (eQTM)). The BIOS-BBMRI database was also used to seek genetic variants influencing methylation levels of nearby or far-away genes (*cis*- and *trans*-methylation quantitative trait loci (meQTL)).

### Functional and regulatory annotation of CpG sites

We conducted hypergeometric tests with Bonferroni correction to compare the genomic characteristics of the replicated CpGs with the whole set of CpGs that we analyzed, using the Infinium Human Methylation 450 Bead-Chip annotation files. Further, we queried cis-meQTLs in the platform of Functional Mapping and Annotation of Genome-Wide Association Studies (FUMA GWAS)^40^. Using this platform, we examined the overlap between cis-meQTLs with signals in the NHGRI-EBI Catalog of published GWAS^41^. As epigenetic signatures are tissue dependent, and our analysis was limited in blood samples, we used the GTEx expression database - which provides an insight into differential expression of relevant genes across different human tissues. For this analysis, we used genes annotated by Illumina 450K (or the nearest gene) to the significantly associated CpGs. Moreover, we searched PubMed using the ‘CpG id’ as the keyword to search any potential links between significant CpGs and different health outcomes. Pathway analysis for the annotated genes was also performed using the IPA software (https://www.qiagenbioinformatics.com/products/ingenuity-pathway-analysis/).

### Mendelian Randomization (MR) study

We implemented a two-sample Mendelian randomization (MR) approach to evaluate the potential causal effect of coffee consumption on the identified CpGs, investigating whether the DNA methylation changes are a consequence of coffee consumption (**Figure S1**). To this end, we used 50 independent SNPs reported in previous GWASs on coffee consumption as instrumental variables (IVs) (**Table S2**)^42,43^.

In addition, we assessed the potential causal association of coffee-related CpGs with a number of cardiovascular and metabolic traits, including coronary heart disease (CHD), type 2 diabetes (T2D), BMI, waist-hip ratio (WHR), lipid traits (HDL-C, LDL-C, total cholesterol, triglycerides), and fatty liver disease. For each CpG, we calculated instrumental variables for DNA methylation levels based on methylation quantitative trait loci (cis-meQTL) obtained from FHS cohort (N~4,170)^44^. Two methods were used to explore causality. First, a weighted genetic risk score (GRS) was constructed for coffee consumption. The other MR approaches implemented were inverse variance weighting (IVW), and sensitivity MR analyses: weighted median and MR-Egger. We used MR-PRESSO (Mendelian Randomization pleiotropy residual sum and outlier) to identify horizontal pleiotropic outliers in multi-instrument summary-level MR testing (https://github.com/rondolab/MR-PRESSO)^45^. All MR methods for multiple genetic instruments were conducted using the statistical “MendelianRandomization” R-package^46^. Additional information of the MR methods implemented in this study is outlined in Supplementary Information.

### Association of coffee consumption, DNA methylation and liver function

Due to the well-documented association between coffee consumption and liver function^14,47^, we also ran a three-way association to assess the correlation of a coffee-associated CpG with liver enzymes and fatty liver disease in the Rotterdam Study (**Figure S2)**. We first tested the cross-sectional associations between coffee consumption and liver enzymes in the Rotterdam Study (n=5,192). Serum GGT, ALT and AST levels were determined using Merck Diagnostica kit on an Elan Autoanalyzer (Merck, Darmstadt, Germany). The liver enzymes were log transformed to obtain normal distribution. Linear regression models were implemented where each liver enzyme was an outcome, and main exposure was coffee consumption (cups/day) adjusted for sex, age, smoking, BMI and excessive alcohol consumption. Excessive alcohol consumption was defined as >14 units/week for women and >21 units/week for men. Next, we tested the association of the coffee-related CpG with liver enzymes in the Rotterdam Study (n=1,406)^14^. Generalized linear mixed effects models were fitted using the R package lme4 and liver enzymes were log transformed to obtain normal distribution. Three models were analyzed, where each liver enzyme was an outcome, adjusted for age, sex, BMI, smoking, whole blood cells proportion, batch effects and excessive alcohol consumption. All analyses were performed using the statistical package R, version 3.0.2.

### Quantitative RT-PCR and knockdown of *PHGDH* in liver cell lines

Seven established human hepatoma cell lines (including PLC/PRF/5, HepG2, HepRG, Hep3B, SNU398, SNU449 and Huh6) were cultured separately. HepG2, Hep3B, SNU398, SNU449 and Huh6 were cultured in Dulbecco’s modified Eagle’s medium (Invitrogen-Gibco, Breda, the Netherlands) complemented with 10% (v/v) fetal calf serum (Hyclone, Lonan, UT), 100 IU/ml penicillin, 100 μg/ml streptomycin, and 2 mM L-glutamine (Invitrogen-Gibco). The hepatoblastoma cell line PLC/PRF/5 was cultured on fibronectin/collagen/albumin-coated plates (AthenaES) in Williams E medium (Invitrogen-Gibco, Breda, the Netherlands) complemented with 10% (v/v) fetal calf serum, 100 IU/ml penicillin, 100 μg/ml streptomycin, and 2 mM L-glutamine. The human liver progenitor cell line - HepaRG was cultured in William’s E medium supplemented with 10% (v/v) fetal calf serum, 100 IU/ml penicillin, 100 μg/ml streptomycin, 5 μg/ml insulin (Sigma-Aldrich, St. Louis, MO), and 50 μM hydrocortisone hemisuccinate (Sigma-Aldrich, St. Louis, MO). The identity of all cell lines was confirmed by STR genotyping.

RNA was isolated using the Machery-NucleoSpin RNA II kit (Bioke, Leiden, The Netherlands) and quantified using a Nanodrop ND-1000 (Wilmington, DE, USA). cDNA was synthesized from total RNA using a cDNA Synthesis Kit (TAKARA BIO INC). The cDNA of all target genes was amplified for 50 cycles and quantified with a SYBRGreen-based real-time PCR (Applied Biosystems) according to the manufacturer’s instructions. *GAPDH* was considered as a reference gene to normalize gene expression. Relative gene expression was normalized to *GAPDH* using the formula 2^-ΔΔCT^ (ΔΔCT = ΔCTsample – ΔCTcontrol). All primer sequences are included in **Table S3**.

Lentiviral pLKO knockdown vectors (Sigma–Aldrich) targeting *PHGDH* or control were obtained from the Erasmus Biomics Center and produced in HEK293T cells. After a pilot study, the shRNA vectors exerting optimal gene knockdown were selected (shRNA sequences: CCGGCAGACTTCACTGGTGT CAGATCTCGAGATCTGACACCAGTGAAGTCTGTTTTT and CCGGCGCAGAACTCACTTGTG GAATCTCGAGATTCCACAAGTGAGTTCTGCGTTTTT, target sequences are CAGACTTCACTG GTGTCAGAT and CGCAGAACTCACTTGTGGAAT, respectively). Stable gene knockdown cells were generated after lentiviral vector transduction and puromycin (2.5μg/ml; Sigma) selection. The relative expression levels of *PHGDH* with nine lipid-associated genes, reported in the previous GWAS and experimental studies^48–50^, were examined.

## Results

Characteristics of the cohorts participating in the discovery phase (n=9,612) and replication phase (n=6,177) are presented in **Table 1**. The mean age across all participating cohorts ranged from 41.1 years in Airwave cohort to 78.6 years in CHS_EA cohort. The majority of the study participants were women (61.44%). Mean total coffee intake among cohorts ranged from 0.6 cups/day in the CHS_AA cohort to 3.5 cups/day in the RS-III-2 cohort, while mean total tea intake ranged from 0.3 cup/day in the EPIC_Italy cohort to 3.4 cups/day in TwinsUK (**Table 1**).

The quantile-quantile (QQ) plots were generated and corresponding lambda value computed for the overall meta-analysis of the discovery and replication panels combined, indicated no statistical inflation in the fully adjusted models for coffee or tea consumption (**Figure S3**). Furthermore, we inspected effect-size distribution plots indicating that one cohort (Lifelines, n=186) had an effect-size scale non-comparable to other participating cohorts (**Figure S4**). As the Lifelines cohort had a high standard deviation for coffee consumption, and also no data on tea consumption, we excluded this cohort from further analysis.

EWAS meta-analysis of 9,612 participants with European ancestry in the discovery phase identified 11 CpG sites associated with coffee consumption at a suggestive significant level (*P* < 5.0×10^-6^) in model 2 (**Table 2A**). We sought for replication of these CpGs in independent cohorts of both ancestries (EA and AA) comprising 6,177 participants, where 7 CpGs were successfully replicated (0.05/11 CpGs *P* < 4.5×10^-3^). In the combined meta-analysis of discovery and replication cohorts, 11 CpGs passed the epigenome-wide significance threshold (*P* < 1.0× 10^-7^) (**Table 2A**). A Manhattan plot showing the EWAS on coffee consumption is depicted in **Figure 2A**. Forest plots for the significantly associated CpGs showed small effects but an overall consistent direction across participating cohorts (**Figure S5**). Heterogeneity was also assessed; for those CpGs showing nominal evidence of heterogeneity (*P* < 0.05), we additionally provided results from random-effects inverse-variance metaanalysis (**Table S4**). The CpG with the most significant association with coffee consumption was cg05575921 (*P*= 2.17×10^-15^, β= −0.0016) annotated to *AHRR,* repressor of the *AHR* (Aryl Hydrocarbon Receptor) gene (**Figure 3A**).

**Figure 2.**
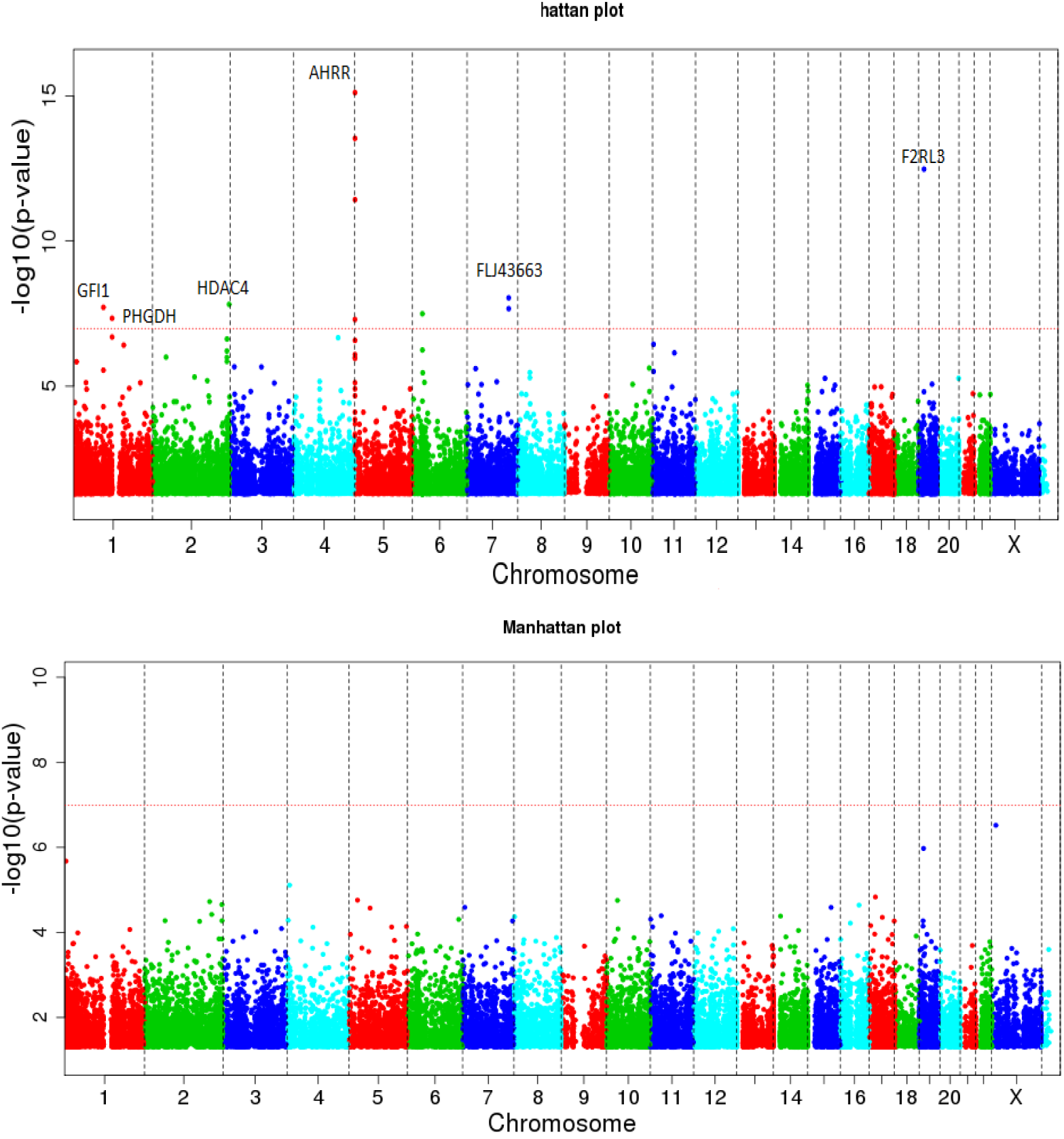
Manhattan plots depicting the results of EWAS meta-analysis in the overall sample for coffee (**A**) and tea (**B**) consumption in the fully adjusted model. The red line indicates the genome-wide significance *P*-value threshold of 1.1×10^-7^.

**Table 2A.**
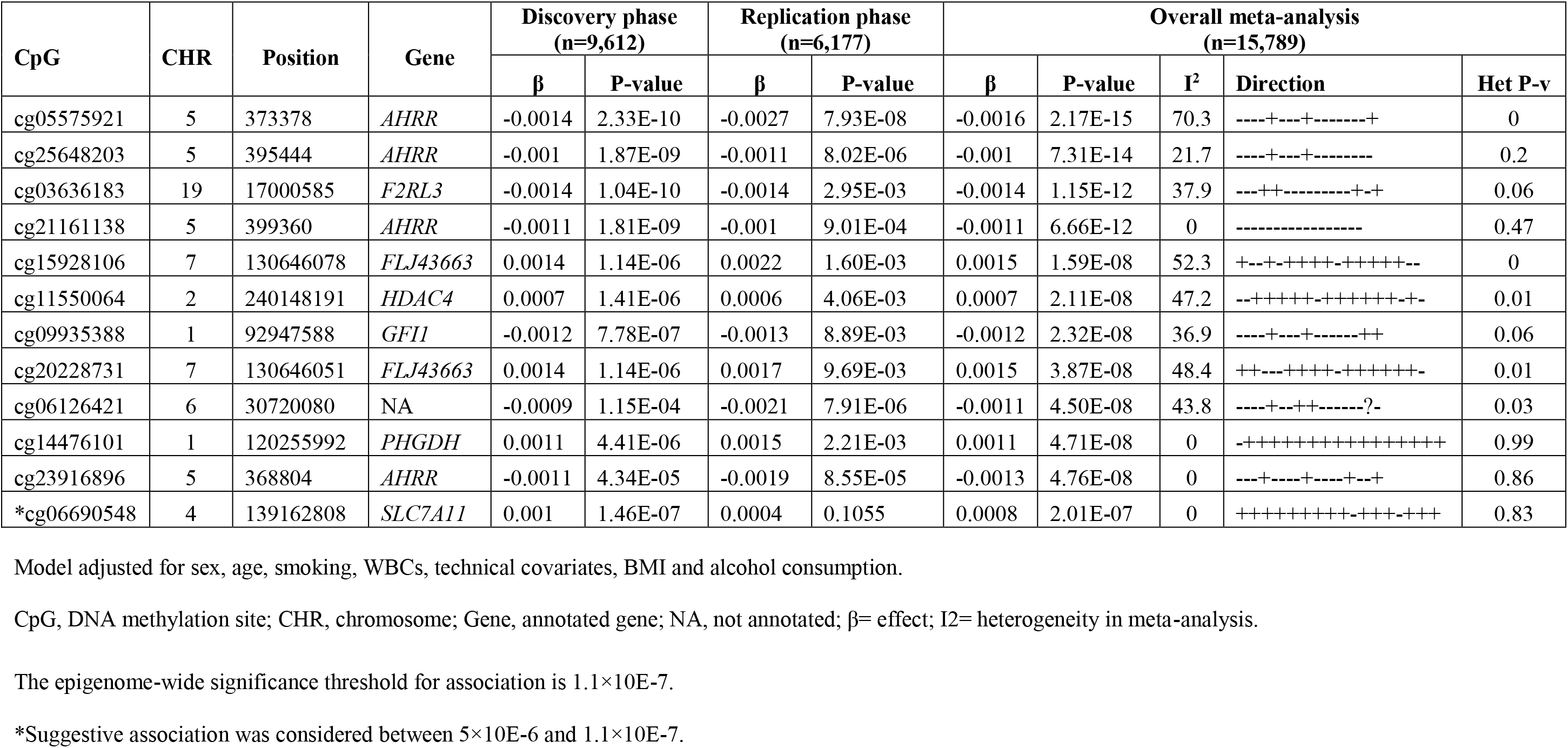
Inverse-variance weighted fixed effects meta-analysis of EWAS with coffee consumption

**Table 2B.**
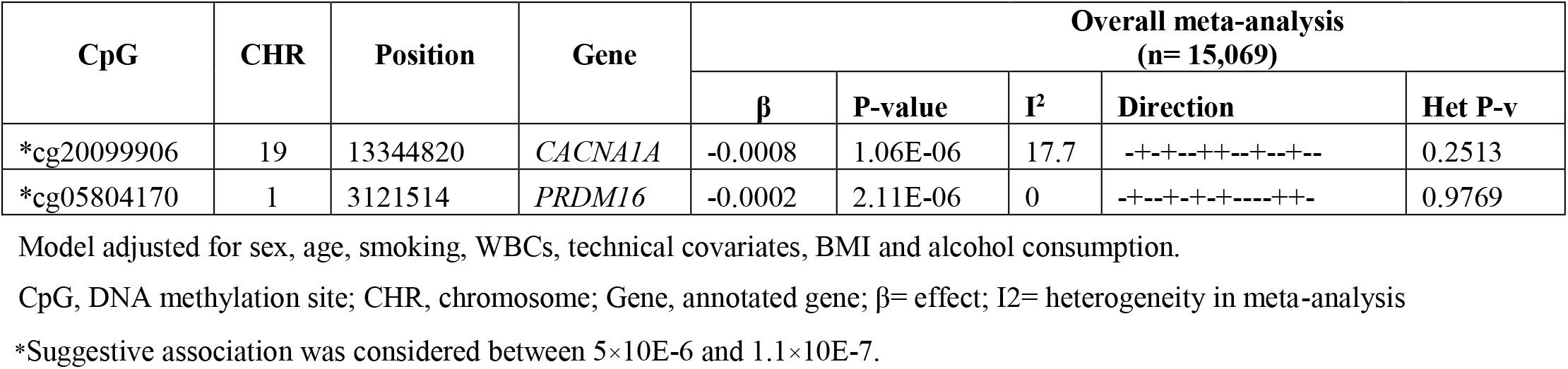
Inverse-variance weighted fixed effects meta-analysis of EWAS with tea consumption

EWAS meta-analysis of tea consumption from all participating cohorts (n=15,789) showed two suggestively associated CpGs (*P* < 5.0×10^-6^), namely cg20099906 *(CACNA1A)* and cg0584170 *(PRDM16)* (**Table 2B**). The Manhattan plot showing the EWAS results on tea consumption is depicted in **Figure 2B**.

When excluding the cohorts with different time points of collection between methylation and beverage intake data, the results of association between DNA methylation levels and coffee, and tea consumptions did not change substantially (**Table S5**). Furthermore, investigating potential ancestry effects, we performed an EWAS meta-analysis of coffee consumption separately in EA (n=12,868) and AA (n=2,921) participants. Our results showed that out of the 9 CpGs significantly associated with coffee consumption in EA participants (n=12,868), none replicated in AA participants (*P* < 0.005 (0.05/9)). Conversely, EWAS in AA participants (n=2,921) showed one CpG site (cg05822739) associated with coffee consumption (*P*= 1.08×10^-7^, β= −0.0015), which was not identified in EA participants despite the larger sample size (**Table S6**).

Of the 11 CpGs significantly associated with coffee consumption, nine have been annotated to the genes *AHRR, F2RL3, FLJ43663, HDAC4, GFI1,* and *PHGDH* **(Figure 3B)**. A heatmap depicting average expression of these 6 genes across 53 human tissues, available on the “Functional Mapping and Annotation of genetic associations with FUMA” webtool^40^, is provided in the **Figure S6**. Based on the tissue specificity of differential expression using FUMA, these genes showed an up-regulation in the transverse colon while they seemed to be down regulated in the coronary artery. Furthermore, esophagus muscularis and lung tissue showed a differential expression of the genes (**Figure S7**). The pathway analysis using IPA for the 6 annotated genes showed enrichment for serine biosynthesis (*P*= 1.36×10^-3^) and xenobiotic metabolism signalling (*P*= 2.71×10^-3^) and association with inflammatory response (*P*-value between 4.48×10^-2^ and 4.42×10^-5^) (**Table S7**).

**Figure 3.**
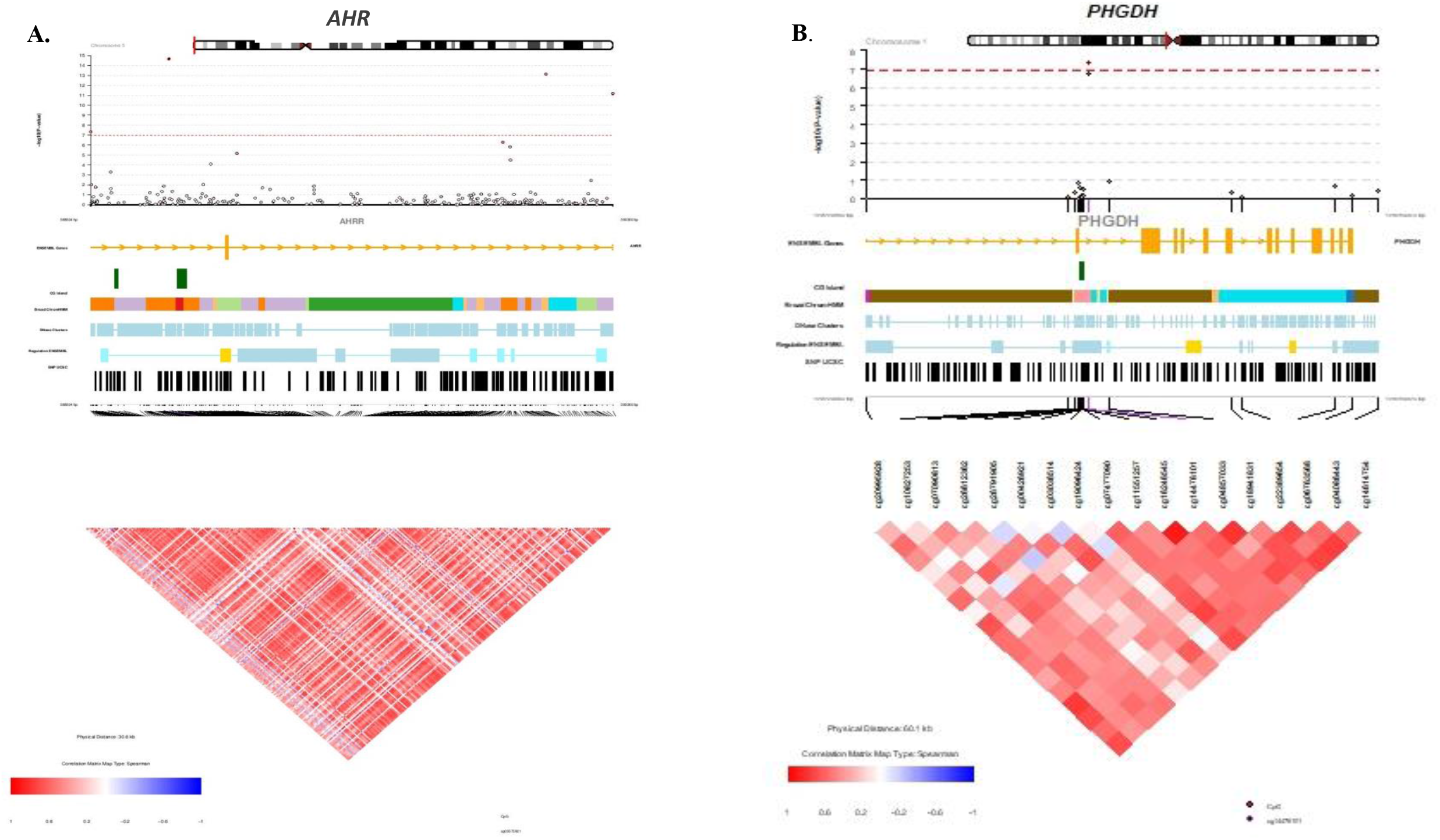
The CoMET plots depicting genomic regions where the CpGs annotated to the *AHRR* gene (**A**) and the *PHGDH* gene (**B**) are located. The x-axis indicates the position in base pair (bp) (hg19) for the region, while y-axis indicates the strength of association from EWAS with coffee consumption. The red line indicates the Bonferroni threshold for Epigenome-wide significance (*P*= 1.1× 10^-7^). The figure was computed using the R-based package CoMET, while the Ensembl is a genome database resource (http://ensemblgenomes.org/). The correlation of the surrounding CpGs was computed using methylation measures in the Rotterdam Study.

The 11 coffee-associated CpGs were further explored in association with genetic variation (meQTL) or expression levels of their nearby or distant genes (eQTM) using the BIOS-BBMRI database. Eight of the CpGs were associated with genetic variants in the neighbouring genes (*cis*-meQTLs) (**Table S8**). By overlapping *cis*-meQTL variants with GWAS results in the NHGRI-EBI GWAS Catalog, we did not find *cis*-meQTLs or their proxies (LD R^2^ > 0.8) to be associated with any traits in previous GWASs. Furthermore, 6 of the 11 CpGs showed association with expression levels of their nearby genes (eQTM) (**Table S8**), including 3 CpGs annotated to *AHRR* that were associated with expression of *EXOC3.* Among the 11 CpGs, the most significant association with eQTM was between cg14476101 and expression levels of *PHGDH (P=* 2.05×10^-55^). Literature search for the association between the 11 identified CpGs and any phenotypes or diseases in PubMed showed overlap with some traits that are shown in **Table S9**, in particular between the CpGs annotated to *AHRR* and smoking.

We next assessed the causal association between coffee consumption and the 11 identified CpGs in the RS and FHS. The weighted GRS-based MR analysis did not support the causal association, which might be due to lack of statistical power. For example, we observed non-significant results between coffee consumption and cg14476101 (GRS-β= −3.42×10^-5^, GRS-P= 0.22). Additionally, our results from the multi-IVs, conventional and sensitivity MR analyses for this CpG also did not show significant evidence for causality (IVW-β= 0.01, IVW-P= 0.1) (**Table S10** and **Figure S8**). We also tested the potential causal association of the coffee-associated CpGs with cardiovascular disease and metabolic traits. Multi-instrument MR analyses showed that cg01940273 could be causally associated with T2D, BMI, WHR, LDL-C and total cholesterol (**Figure S9**); cg05575921 with BMI, WHR and HDL-C (**Figure S10**); cg09935388 with T2D and HDL-C (**Figure S11**); cg11550064 with BMI, WHR, HDL-C, LDL-C, total cholesterol, triglycerides and CHD (**Figure S12**); and cg23916896 with T2D, BMI, HDL-C and total cholesterol (**Figure S13**) (**Table S11**). The causal association between cg14476101 and fatty liver disease has been previously confirmed by MR analysis in FHS, where hypermethylation at the locus was associated with lower fatty liver (*P*= 0.01)^51^.

The inverse association of coffee consumption with liver diseases has been well documented by different researchers^14,47,52^. The CpG cg14476101 and its annotated gene *(PHGDH)* have been reported in previous studies to be associated with fatty liver disease and adiposity^53,54^. Methylation-gene expression association between cg14476101 and *PHGDH* has also been verified in liver tissue^54^. Moreover, the expression level of *PHGDH* gene has been associated with liver fat^51^. Thus, we sought to show the three-way association among coffee consumption, DNA methylation of cg14476101 and liver function in the Rotterdam Study. To this end, we tested the association of coffee consumption and three liver enzymes (n=4,756) adjusted for potential confounders, which showed a negative association between coffee consumption and serum levels of AST (*P*= 0.008, β= −0.005) and GGT (*P*= 0.004, β= −0.011) (**Table S12**). In addition, we tested the association of cg14476101 with the liver enzymes (n=1,406) adjusted for age, sex, BMI, smoking, whole blood cells proportion, batch effects and excessive alcohol consumption, and observed a nominal association with the serum levels of AST (*P*= 0.016, β= −0.26) and a suggestive association with GGT (*P*= 0.06, β= −0.43). These data may suggest that the observed link between *PHGDH* expression and fatty liver disease is mediated by coffee consumption via altering DNA methylation levels of cg14476101.

To gain further insight into the mechanism linking *PHGDH* to fatty liver disease, we conducted additional experimental studies. We measured the expression level of *PHGDH* in several human liver cell lines and, subsequently, related it to the expression levels of a panel of lipid-associated genes. **Figure 4A** displays the *PHGDH* expression level in seven liver cell lines. From this, we selected snu398 cells, with the highest expression levels of *PHGDH,* and snu449 cells, with the lowest expression levels of *PHGDH,* and compared the relative expression levels of *PHGDH* with nine know lipid-related genes, reported in the previous GWAS and experimental studies to be involved in lipid metabolism^48–50^. The *PHGDH* expression level was correlated with the expression levels of five of these lipid-associated genes (**Figure 4B**). Next, we knocked down *PHGDH* expression in PLC/PRF/5 cells using lentiviral shRNA vectors (**Figure 4C**). After silencing *PHGDH*, we observed a significant decrease of *LPL* expression and a significant increase of *LDR* and *ABCA1* expression (*P* < 0.05) in both knocked down cells (**Figure 4D**), in line with the observed correlations in snu398 and snu449 cells. These experiments may suggest a potential role of *PHGDH* in lipid metabolism and fat accumulation in the liver, that could occur through regulating the expression of lipid-associated genes.

**Figure 4.**
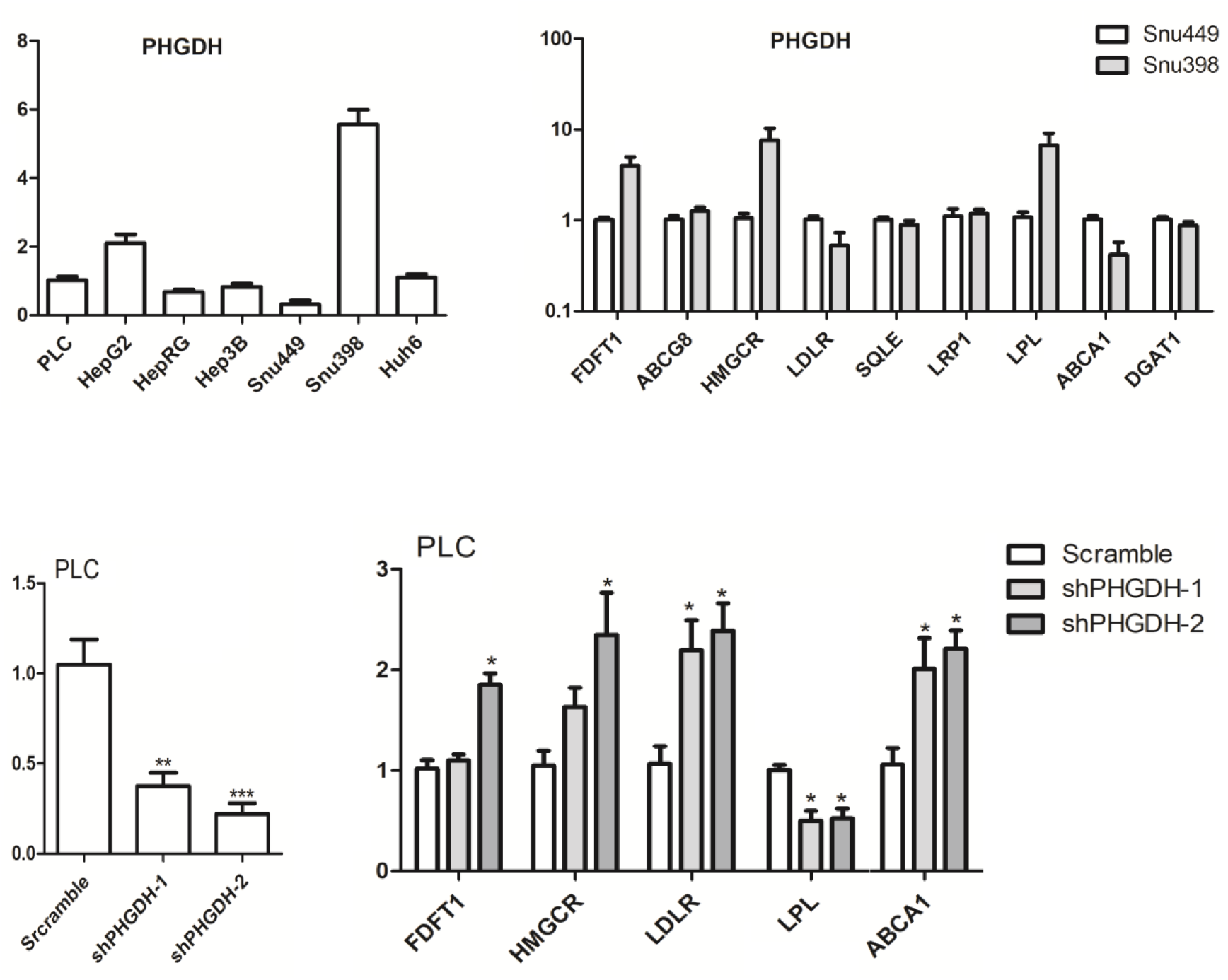
**A**) Relative expression levels of *PHGDH* gene against a reference gene (GAPDH) in 7 human liver cell lines. Gene expression levels were quantified by qRT-PCR. Data were normalized to the PLC cell line (PLC, set as 1). The results are presented as mean ± SEM of three independent experiments with 2-4 biological repeats each. **B**) Relative expression levels of 9 lipid- associated genes in snu499 cell line (with the lowest level of *PHGDH* expression) and snu398 cell line (with the highest level of *PHGDH* expression) are shown. Relative gene expression levels were quantified by qRT-PCR. GAPDH serves as a reference gene, and gene expression levels in snu449 cell line set as 1. Data are means ± SEM of three independent experiments with 2-4 biological repeats each. This figure shows that, compared with snu4499 cells, snu398 cells differentially express five of the lipid-associated genes *(FDFT1, HMGCR, LDLR, LPL,* and *ABCA1*). **C**) Established *PHGHD* knockdown cell lines (shPHGHD-1 and −2), PLC cells transduced with lentiviral shRNA vectors targeting *PHGDH* or scramble control. qRT-PCR analysis of *PHGDH* expression were performed in stable knockdown or scramble control PLC cells. Data are normalized to the scramble control (scramble, set as 1). Data are means ± SEM of three independent experiments with 2-4 biological repeats each, \*\**P*<0.01 and \*\*\**P*<0.001. **D**) Expression levels of five lipid-associated genes in stable *PHGDH* knockdown or scramble control PLC cells. Data were normalized to the scramble control (scramble, set as 1). Data are means ± SEM of three independent experiments with 2-4 biological repeats each. Comparisons between groups were performed with Mann-Whitney test. Differences were considered significant at *P* < 0.05 (indicated by*). The figure demonstrates that knockdown of *PHGDH* gene expression by lentiviral shRNA vectors resulted in significant decrease in the expression level of *LPL* and significant increase in the expression levels of *LDLR* and *ABCA1* in both knockdown cells.

## Discussion

In this study, we conducted the largest EWAS meta-analyses of coffee and tea consumption to date comprising more than 15,000 participants. We found that coffee consumption was associated with differential methylation of peripheral blood-derived DNA at 11 CpG sites. The genes annotated to some of the identified CpGs have potential relevance in pathways underlying coffee metabolism and have been associated with various health outcomes^55–57^. Among these, cg14476101 was significantly associated with expression level of its annotated gene *PHGDH* and a reduced risk of fatty liver disease. Our experimental studies further demonstrated a correlation between expression levels of *PHGDH* and some lipid-associated genes in liver cells, suggesting a potential role of *PHGDH* in hepatic-lipid metabolism.

Our EWAS meta-analysis revealed 11 CpGs significantly associated with coffee consumption, seven of which were negatively associated with coffee intake. Four of these seven CpGs are annotated to the *AHRR* (Aryl-Hydrocarbon Receptor Repressor). This gene encodes a repressor of *AHR,* which has been previously associated with coffee consumption in a large-scale GWAS (n=91,462)^33^. *AHRR* and *AHR,* alongside with *CYP1A1* and *CYP1A2,* belong to the xenobiotic metabolism pathway, in which *AHR* is activated via various ligands such as polycyclic aromatic hydrocarbons (PAHs) or with ligands of natural origins (e.g., food)^58^. PAHs are carcinogenic substances found in tobacco smoke^59^, which can also be formed during coffee roasting processes^60^. It has been established that PAHs are mediating their impact on the cell via *AHR*^61^. Nevertheless, the *AHRR* CpGs identified in our analysis associated with coffee have been also reported in previous EWAS of smoking^62,63^. Although we accounted for smoking status in our analyses, given that cigarette smoking is associated with coffee consumption^64^ and smoking has a notable effect on DNA methylation^62^, the association between these CpGs with coffee consumption might warrant cautious interpretation. We speculate that methylation within *AHRR* might influence the xenobiotic pathway and the subsequent genes *(AHR, CYP1A1* and *CYP2A2)* identified in the coffee consumption GWAS^33^. The other three CpGs negatively associated with coffee consumption in our study are annotated to the *F2RL3, GFI1* and *IER3* genes, which were previously shown to be involved in a wide range of phenotypes from inflammatory response^65,66^, to cancer^67,68^ and cardio-metabolic diseases^69,70^. Because of the strong association between coffee and cardiovascular disease^71^ as well as cancer^72^, further experimental studies are warranted to investigate whether alteration of DNA methylation level at the associated CpGs change gene expression and confer risk for these complex diseases.

Five of the 11 identified CpGs were positively associated with coffee consumption. Four of these are annotated to *FLJ43663, HDAC4* and *PHGDH.* To date, little research has focused on *FLJ43663* in disease predisposition. A recent study showed the involvement of *FLJ43663* as a risk factor for Behçet’s disease^73^ and another study reported that *FLJ43663* gene polymorphisms were associated with the risk of breast cancer in a Han Chinese population^74^. The second gene, *HDAC4* is a member of the xenobiotic metabolism signalling. This gene is involved in cocaine-related behaviours^75^, with a proposed mechanism that cocaine-induced nuclear export of *HDAC4* could promote development of cocaine reward behaviours. Furthermore, animal studies have previously shown that *HDAC4* inhibition increases sensitivity to cocaine, whereas overexpression of the gene has opposite effect, which further supports the importance of *HDAC4* in conditioned place preference in addictive behaviour^76^. Our study is the first to show a link between coffee consumption and *FLJ43663* and *HDAC4* genes, although further investigation is needed to address the exact mechanisms through which these genes might play a role in human diseases.

Finally, *PHGDH* gene is of particular interest. This gene encodes the phosphoglycerate dehydrogenase enzyme that catalyses the first and rate-limiting step in the phosphorylated pathway of serine biosynthesis. Methylation-gene expression association between cg14476101 and *PHGDH* has been demonstrated in blood and liver^54^. The CpG site has been reported to be negatively associated with the levels of liver enzymes in serum^53^ and the risk of non-alcoholic fatty liver disease (NAFLD)^51^. Methylation of cg14476101 has also been linked to adiposity measured by BMI and waist circumference^54^. Another EWAS has reported the association between this CpG and blood concentration of steroid hormones, which are upregulated in obesity^77^. In line with our findings in the Rotterdam Study, several studies have shown an inverse association between coffee consumption and liver enzymes, including ALT, AST, and GGT^78^. Previous studies have also associated coffee consumption with reduced risk of chronic liver disease^47^, hepatocellular carcinoma^79^, and cirrhosis^80^. These findings might proposed a link between *PHGDH* expression and liver function, which can be mediated by coffee consumption via altering the DNA methylation levels of cg14476101. We attempted to provide additional evidence of directionality of coffee consumption on cg14476101 using Mendelian randomization techniques, but the power of our MR analyses was limited due to the lack of availability of genetic data for some cohorts limiting sample size combined with a lack of strong genetic instruments for this CpG. The non-significant results can also be explained by other reasons. Firstly, population stratification, although we used genetic information from large GWAS performed mainly in European population and adjusted by population substructure^43^. Secondly, pleiotropy, since some of the SNPs used as instrumental variables have also been associated with lipid traits and body size which might influence the causal estimates. Yet, the results of MR excluding potential pleiotropic variants were fairly similar and MR-Egger, implemented in this study, is an useful approach to account for pleiotropy^81^. Thirdly, the genetic variants for coffee consumption were associated with number of cups of coffee per day among coffee drinkers, and the effect estimates might not relate to DNA methylation observed among ever/never coffee drinkers^82^.

Furthermore, we knocked down the expression of *PHGDH* in human liver cells and revealed a correlation between expression of this gene and lipid-associated genes (*LPL*, *LDLR* and *ABCA1*), which suggest a potential role of *PHGDH* in hepatic-lipid metabolism. Previous evidence has also indicated that reduced expression of *PHGDH* is closely associated with the development of fatty liver disease^57^. In addition, our results showed a suggestive association between coffee consumption and cg06690548 annotated to *SLC7A11* (β=0.0008, *P*= 2.0×10^-7^). The *SLC7A11* gene is known as transporter of cysteine and glutamate, whereas caffeine promotes glutamate release in the posterior hypothalamus^83^. The CpG has been also reported in two EWASs to be associated with liver enzymes and NAFLD^51,53^. Moreover, similar experimental studies demonstrated the involvement of *SLC7A11* in hepatic-lipid metabolism^53^. Thus, more experimental studies are merited to further elucidate in what way epigenetic modification of *PHGDH* and *SLC7A11* could explain the beneficial effect of coffee consumption on liver diseases.

In our study, EWAS meta-analysis of tea consumption showed only two suggestive associations, despite having the advantage of much larger sample size and good statistical power compared to the previously published EWA study^20^. In the earlier study, Ek *et al.* reported two CpGs associated with tea consumption in women, nevertheless, those CpGs were not replicated in our study. Lack of statistically significant association between tea consumption compared with coffee in our analysis can be explained by different bioavailability of polyphenols and other chemical compounds or much lower concentration of caffeine in tea compared to coffee. Among the two CpGs suggestively associated with tea consumption, cg20099906 is annotated to *CANA1A* (Calcium Voltage-Gated Channel Subunit Alpha1 I). It has previously been shown that food polyphenols can inhibit cardiac voltage-gated sodium channels^84^. In addition, the major compound of tea is a polyphenol called epigallocatechin-3-gallate, that was shown in an *in vivo* study to modulate the Ca^2+^ signals^85^. This might indicate a potential link between DNA methylation of cg20099906 and cardiovascular diseases. Future studies with even larger sample sizes are needed to replicate these suggestive associations with tea intake and investigate their potential links with different health outcomes.

The major strength of the present study is the large sample size and multi-ethnic contribution. All contributing cohorts had DNA methylation measured in whole blood, and adjustment for cell components allowed us to account for different epigenetic markers within cells present in the blood. Incorporation of different adjustment models allowed us to limit confounding to a certain extent. Yet, the findings of this study should be considered in light of some limitations. One important concern regarding this analysis is smoking, given that the effect of smoking on DNA methylation has been recognized^62^ and previous studies have shown that heavier smokers tend to drink more coffee^64^. Even though we adjusted for smoking in our analysis, some differences might not be tackled by our statistical approach and might yield confounded analysis to some extent. Also, we cannot rule out the possibility of residual confounding in our analysis, given that some of the participating cohorts did not have data for current, former and never smokers. Furthermore, smoking status might be misclassified or the possibility of second-hand smoke cannot be ruled out. The results presented could reflect potential pleiotropy, confounding or both, or it could provide insight into the potential causal role of coffee with DNA methylation and disentangling these would merit further investigations. In addition, only total coffee was assessed and was used as a continuous variable (cups/day), and some cohorts have different cup sizes. However, we believe that this limitation would rather dilute the findings. We also did not include information on coffee brewing methods, which might have a large effect on what compounds are in the final beverage (e.g. filtered *vs* non-filtered coffee). Finally, since our study consists mainly of middle aged and elderly individuals of two ancestries, other studies are needed to assess the generalizability of our findings to other age groups and ancestries.

The present study is thus far the largest EWAS meta-analyses exploring the association of coffee and tea consumption with DNA methylation. We found that coffee consumption is significantly associated with differential DNA methylation at 11 CpGs in blood. The genes annotated to some of these CpGs have potential relevance in pathways underlying coffee metabolism and have been linked to risk of diseases. These findings may provide new insights into the mechanism of action of coffee consumption in conferring disease risk. Future studies with larger and more ethnically diverse sample sizes are warranted to validate our findings and to explore the biological relevance of the associated DNA methylation sites and genes in beneficial and harmful association with different health outcomes.

## Data availability

The datasets generated during this study are available from the corresponding author upon reasonable request.

## Acknowledgments

The authors are grateful to the staff and participants of all cohorts involved in this study for their important contributions. Specific funding and acknowledgements statements of each study can be found in Supplementary Information.

## Author contributions

MG, IK, and MK contributed to study design. IK, EP, JM, SM, DS, BK, YZ, SA, GF, JH, JEC, NK, BS, and SAG contributed to cohort-specific data analyses. IK contributed to meta-analyses of EWAS. EP and JM contributed to mQTL and Mendelian Randomization analyses. MG, YL and QP designed and performed experimental studies in liver cell lines. TV, KB, MAI, JLT, EAH, CC, PCT, RC, JM, AGU, MW, DL, MK, MF, MW, HB, ZH, CR, PV, PE, JTB, NS, AP, AD, LF, SP, NV, VC, SG, and RJK contributed to cohort design and management, and data collection. MG, IK, and EP contributed to interpretation of the results and writing of manuscript. MG, IK, EP, SM, MK, JH, CMR, DL, BS, HB, JLT, KW, MW, SA, SG, AK contributed to critical review of manuscript.

## References

1 Landais, E. et al. Coffee and tea consumption and the contribution of their added ingredients to total energy and nutrient intakes in 10 European countries: Benchmark data from the late 1990s. Nutrients 10, 725 (2018).

2 National Research Council Committee on, D. & Health. doi:NBK218743. 10.17226/1222 (1989).

3 Temple, J. L. et al. The Safety of Ingested Caffeine: A Comprehensive Review. Front Psychiatry 8, 80, doi:10.3389/fpsyt.2017.00080 (2017).

4 Bunker, M. L. & McWilliams, M. Caffeine content of common beverages. J Am Diet Assoc 74, 28–32 (1979).

5 Rein, M. J. et al. Bioavailability of bioactive food compounds: a challenging journey to bioefficacy. Br J Clin Pharmacol 75, 588–602, doi:10.1111/j.1365-2125.2012.04425.x (2013).

6 Ferruzzi, M. G. The influence of beverage composition on delivery of phenolic compounds from coffee and tea. Physiol Behav 100, 33–41, doi:10.1016/j.physbeh.2010.01.035 (2010).

7 Fuller, M. & Rao, N. Z. The Effect of Time, Roasting Temperature, and Grind Size on Caffeine and Chlorogenic Acid Concentrations in Cold Brew Coffee. Sci Rep-Uk 7, doi:ARTN 17979. 10.1038/s41598-017-18247-4 (2017).

8 Nieber, K. The Impact of Coffee on Health. Planta Med 83, 1256–1263, doi:10.1055/s-0043-115007 (2017).

9 Khan, N. & Mukhtar, H. Tea and health: studies in humans. Curr Pharm Des 19, 6141–6147, doi:CPD-EPUB-20130219-6 (2013).

10 van Dieren, S. et al. Coffee and tea consumption and risk of type 2 diabetes. Diabetologia 52, 2561–2569, doi:10.1007/s00125-009-1516-3 (2009).

11 Bohn, S. K., Ward, N. C., Hodgson, J. M. & Croft, K. D. Effects of tea and coffee on cardiovascular disease risk. Food Funct 3, 575–591, doi:10.1039/c2fo10288a (2012).

12 Jee, S. H. et al. Coffee consumption and serum lipids: a meta-analysis of randomized controlled clinical trials. American journal of epidemiology 153, 353–362 (2001).

13 Loftfield, E. et al. Association of Coffee Drinking With Mortality by Genetic Variation in Caffeine Metabolism: Findings From the UK Biobank. JAMA Intern Med 178, 1086–1097, doi:2686145.10.1001/jamainternmed.2018.2425 (2018).

14 Alferink, L. J. M. et al. Coffee and herbal tea consumption is associated with lower liver stiffness in the general population: The Rotterdam study. J Hepatol 67, 339–348, doi:S0168-8278(17)30147-2.10.1016/j.jhep.2017.03.013 (2017).

15 Heath, R. D., Brahmbhatt, M., Tahan, A. C., Ibdah, J. A. & Tahan, V. Coffee: The magical bean for liver diseases. World J Hepatol 9, 689–696, doi:10.4254/wjh.v9.i15.689 (2017).

16 Hamilton, J. P. Epigenetics: principles and practice. Dig Dis 29, 130–135, doi:000323874.10.1159/000323874 (2011).

17 Alegria-Torres, J. A., Baccarelli, A. & Bollati, V. Epigenetics and lifestyle. Epigenomics 3, 267–277, doi:10.2217/epi.11.22 (2011).

18 Weinhold, B. Epigenetics: the science of change. Environ Health Perspect 114, A160–167, doi:10.1289/ehp.114-a160 (2006).

19 Anderson, O. S., Sant, K. E. & Dolinoy, D. C. Nutrition and epigenetics: an interplay of dietary methyl donors, one-carbon metabolism and DNA methylation. J Nutr Biochem 23, 853–859, doi:S0955-2863(12)00059-9.10.1016/j.jnutbio.2012.03.003 (2012).

20 Ek, W. E. et al. Tea and coffee consumption in relation to DNA methylation in four European cohorts. Hum Mol Genet 26, 3221–3231, doi:3848993 (2017).

21 Chuang, Y. H. et al. Coffee consumption is associated with DNA methylation levels of human blood. Eur J Hum Genet 25, 608–616, doi:10.1038/ejhg.2016.175 (2017).

22 Psaty, B. M. et al. Cohorts for Heart and Aging Research in Genomic Epidemiology (CHARGE) Consortium Design of Prospective Meta-Analyses of Genome-Wide Association Studies From 5 Cohorts. Circ-Cardiovasc Gene 2, 73–U128, doi:10.1161/Circgenetics.108.829747 (2009).

23 Elliott, P. et al. The Airwave Health Monitoring Study of police officers and staff in Great Britain: rationale, design and methods. Environmental Research 134, 280–285 (2014).

24 Fraser, A. et al. Cohort Profile: The Avon Longitudinal Study of Parents and Children: ALSPAC mothers cohort. International Journal of Epidemiology 42, 97–110, doi:10.1093/ije/dys066 (2013).

25 Low, M., Stegmaier, C., Ziegler, H., Rothenbacher, D. & Brenner, H. Epidemiological investigations of the chances of preventing, recognizing early and optimally treating chronic diseases in an elderly population (ESTHER study). Deut Med Wochenschr 129, 2643–2647, doi:10.1055/s-2004-836089 (2004).

26 Kannel, W. B., Feinleib, M., Mcnamara, P. M., Garrison, R. J. & Castelli, W. P. Investigation of Coronary Heart-Disease in Families - Framingham Offspring Study. American Journal of Epidemiology 110, 281–290, doi:DOI 10.1093/oxfordjournals.aje.a112813 (1979).

27 Wichmann, H. E., Gieger, C., Illig, T. & Grp, M. K. S. KORA-gen - Resource for population genetics, controls and a broad spectrum of disease phenotypes. Gesundheitswesen 67, S26–S30, doi:10.1055/s-2005-858226 (2005).

28 Ikram, M. A. et al. The Rotterdam Study: 2018 update on objectives, design and main results. European journal of epidemiology 32, 807–850 (2017).

29 Moayyeri, A., Hammond, C. J., Hart, D. J. & Spector, T. D. The UK adult twin registry (TwinsUK Resource). Twin Research and Human Genetics 16, 144–149 (2013).

30 Aric, I. The atherosclerosis risk in communit (aric) stui) y: Design and objectwes. American journal of epidemiology 129, 687–702 (1989).

31 Fried, L. P. et al. The cardiovascular health study: design and rationale. Annals of epidemiology 1, 263–276 (1991).

32 Bingham, S. & Riboli, E. Diet and cancer—the European prospective investigation into cancer and nutrition. Nature Reviews Cancer 4, 206–215 (2004).

33 Coffee et al. Genome-wide meta-analysis identifies six novel loci associated with habitual coffee consumption. Mol Psychiatry 20, 647–656, doi:10.1038/mp.2014.107 (2015).

34 Yang, T. O., Crowe, F., Cairns, B. J., Reeves, G. K. & Beral, V. Tea and coffee and risk of endometrial cancer: cohort study and meta-analysis. Am J Clin Nutr 101, 570–578, doi:10.3945/ajcn.113.081836 (2015).

35 Sandoval, J. et al. Validation of a DNA methylation microarray for 450,000 CpG sites in the human genome. Epigenetics 6, 692–702 (2011).

36 Houseman, E. A. et al. DNA methylation arrays as surrogate measures of cell mixture distribution. BMC bioinformatics 13, 86 (2012).

37 Van der Most, P. J., Kupers, L. K., Snieder, H. & Nolte, I. QCEWAS: automated quality control of results of epigenome-wide association studies. Bioinformatics 33, 1243–1245, doi:10.1093/bioinformatics/btw766 (2017).

38 Willer, C. J., Li, Y. & Abecasis, G. R. METAL: fast and efficient meta-analysis of genomewide association scans. Bioinformatics 26, 2190–2191 (2010).

39 Magi, R. & Morris, A. P. GWAMA: software for genome-wide association meta-analysis. BMC Bioinformatics 11, 288, doi:1471-2105-11-288 (2010).

40 Watanabe, K., Taskesen, E., van Bochoven, A. & Posthuma, D. Functional mapping and annotation of genetic associations with FUMA. Nat Commun 8, 1826, doi:10.1038/s41467-017-01261-5 (2017).

41 Galperin, M. Y., Fernandez-Suarez, X. M. & Rigden, D. J. The 24th annual Nucleic Acids Research database issue: a look back and upcoming changes. Nucleic Acids Res 45, D1–D11, doi:10.1093/nar/gkw1188 (2017).

42 Kwok, M. K., Leung, G. M. & Schooling, C. M. Habitual coffee consumption and risk of type 2 diabetes, ischemic heart disease, depression and Alzheimer’s disease: a Mendelian randomization study. Sci Rep-Uk 6, 36500 (2016).

43 Kennedy, O. J. et al. Coffee Consumption and Kidney Function: A Mendelian Randomization Study. American Journal of Kidney Diseases (2019).

44 Huan, T. et al. Genome-wide identification of DNA methylation QTLs in whole blood highlights pathways for cardiovascular disease. Nature communications 10, 1–14 (2019).

45 Verbanck, M., Chen, C.-y., Neale, B. & Do, R. Detection of widespread horizontal pleiotropy in causal relationships inferred from Mendelian randomization between complex traits and diseases. Nature genetics 50, 693–698 (2018).

46 Yavorska, O. O. & Burgess, S. MendelianRandomization: an R package for performing Mendelian randomization analyses using summarized data. International journal of epidemiology 46, 1734–1739 (2017).

47 Ruhl, C. E. & Everhart, J. E. Coffee and tea consumption are associated with a lower incidence of chronic liver disease in the United States. Gastroenterology 129, 1928–1936, doi:S0016-5085(05)01774-9.10.1053/j.gastro.2005.08.056 (2005).

48 Tabassum, R. et al. Genetic architecture of human plasma lipidome and its link to cardiovascular disease. Nat Commun 10, 4329, doi:10.1038/s41467-019-11954-8 10.1038/s41467-019-11954-8 [pii] (2019).

49 Willer, C. J. et al. Discovery and refinement of loci associated with lipid levels. Nat Genet 45, 1274–1283, doi:10.1038/ng.2797 (2013).

50 Kathiresan, S. et al. Common variants at 30 loci contribute to polygenic dyslipidemia. Nat Genet 41, 56–65, doi:10.1038/ng.291 (2009).

51 Ma, J. et al. A Peripheral Blood DNA Methylation Signature of Hepatic Fat Reveals a Potential Causal Pathway for Non-Alcoholic Fatty Liver Disease. Diabetes, doi:db18-1193.10.2337/db18-1193 (2019).

52 Saab, S., Mallam, D., Cox, G. A., 2nd & Tong, M. J. Impact of coffee on liver diseases: a systematic review. Liver Int 34, 495–504, doi:10.1111/liv.12304 (2014).

53 Nano, J. et al. Epigenome-Wide Association Study Identifies Methylation Sites Associated With Liver Enzymes and Hepatic Steatosis. Gastroenterology 153, 1096–+, doi:10.1053/j.gastro.2017.06.003 (2017).

54 Wahl, S. et al. Epigenome-wide association study of body mass index, and the adverse outcomes of adiposity. Nature 541, 81–86, doi:nature20784.10.1038/nature20784 (2017).

55 Zhang, Y. et al. F2RL3 methylation in blood DNA is a strong predictor of mortality. International journal of epidemiology 43, 1215–1225 (2014).

56 Cole, J. W. & Xu, H. (Am Heart Assoc, 2015).

57 Sim, W.-C. et al. Downregulation of PHGDH expression and hepatic serine level contribute to the development of fatty liver disease. Metabolism 102, 154000 (2020).

58 Larigot, L., Juricek, L., Dairou, J. & Coumoul, X. AhR signaling pathways and regulatory functions. Biochim Open 7, 1–9, doi:10.1016/j.biopen.2018.05.001 (2018).

59 Vu, A. T. et al. Polycyclic Aromatic Hydrocarbons in the Mainstream Smoke of Popular U.S. Cigarettes. Chem Res Toxicol 28, 1616–1626, doi:10.1021/acs.chemrestox.5b00190 (2015).

60 Houessou, J. K. et al. Effect of roasting conditions on the polycyclic aromatic hydrocarbon content in ground Arabica coffee and coffee brew. J Agric Food Chem 55, 9719–9726, doi:10.1021/jf071745s (2007).

61 Chavan, H. & Krishnamurthy, P. Polycyclic aromatic hydrocarbons (PAHs) mediate transcriptional activation of the ATP binding cassette transporter ABCB6 gene via the aryl hydrocarbon receptor (AhR). J Biol Chem 287, 32054–32068, doi:10.1074/jbc.M112.371476 (2012).

62 Joehanes, R. et al. Epigenetic Signatures of Cigarette Smoking. Circ Cardiovasc Genet 9, 436–447, doi:10.1161/CIRCGENETICS.116.001506 (2016).

63 Philibert, R. A., Beach, S. R. H., Lei, M.-K. & Brody, G. H. Changes in DNA methylation at the aryl hydrocarbon receptor repressor may be a new biomarker for smoking. Clinical epigenetics 5, 19 (2013).

64 Bjorngaard, J. H. et al. Heavier smoking increases coffee consumption: findings from a Mendelian randomization analysis. Int J Epidemiol 46, 1958–1967, doi:4082629 (2017).

65 Fu, Q. et al. Protease-activated receptor 4: a critical participator in inflammatory response. Inflammation 38, 886–895 (2015).

66 Arlt, A. & Schäfer, H. Role of the immediate early response 3 (IER3) gene in cellular stress response, inflammation and tumorigenesis. European journal of cell biology 90, 545–552 (2011).

67 Gratio, V., Walker, F., Lehy, T., Laburthe, M. & Darmoul, D. Aberrant expression of proteinase-activated receptor 4 promotes colon cancer cell proliferation through a persistent signaling that involves Src and ErbB-2 kinase. International journal of cancer 124, 1517–1525 (2009).

68 Khandanpour, C. et al. A variant allele of Growth Factor Independence 1 (GFI1) is associated with acute myeloid leukemia. Blood, The Journal of the American Society of Hematology 115, 2462–2472 (2010).

69 Leger, A. J., Covic, L. & Kuliopulos, A. Protease-activated receptors in cardiovascular diseases. Circulation 114, 1070–1077 (2006).

70 Parmar, P. et al. Association of maternal prenatal smoking GFI1-locus and cardio-metabolic phenotypes in 18,212 adults. EBioMedicine 38, 206–216 (2018).

71 Ding, M., Bhupathiraju, S. N., Satija, A., van Dam, R. M. & Hu, F. B. Long-term coffee consumption and risk of cardiovascular disease: a systematic review and a dose-response meta-analysis of prospective cohort studies. Circulation 129, 643–659, doi:CIRCULATIONAHA.113.005925 (2014).

72 Arab, L. Epidemiologic evidence on coffee and cancer. Nutr Cancer 62, 271–283, doi:920642504 (2010).

73 Yu, H. et al. Epigenome-wide association study identifies Behcet’s disease-associated methylation loci in Han Chinese. Rheumatology 58, 1574–1584 (2019).

74 Xia, P. et al. Polymorphisms in ESR1 and FLJ43663 are associated with breast cancer risk in the Han population. Tumor Biology 35, 2187–2190 (2014).

75 Penrod, R. D. et al. Novel role and regulation of HDAC4 in cocaine-related behaviors. Addict Biol 23, 653–664, doi:10.1111/adb.12522 (2018).

76 Kumar, A. et al. Chromatin remodeling is a key mechanism underlying cocaine-induced plasticity in striatum. Neuron 48, 303–314 (2005).

77 Petersen, A. K. et al. Epigenetics meets metabolomics: an epigenome-wide association study with blood serum metabolic traits. Hum Mol Genet 23, 534–545, doi:10.1093/hmg/ddt430 (2014).

78 Casiglia, E., Spolaore, P., Inocchio, G. & Ambrosio, B. Unexpected effects of coffee consumption on liver enzymes. European journal of epidemiology 9, 293–297 (1993).

79 Bravi, F., Bosetti, C., Tavani, A., Gallus, S. & La Vecchia, C. Coffee reduces risk for hepatocellular carcinoma: an updated meta-analysis. Clinical gastroenterology and hepatology 11, 1413–1421. e1411 (2013).

80 Liu, F. et al. Coffee consumption decreases risks for hepatic fibrosis and cirrhosis: a metaanalysis. PloS one 10, e0142457 (2015).

81 Bowden, J., Davey Smith, G. & Burgess, S. Mendelian randomization with invalid instruments: effect estimation and bias detection through Egger regression. Int J Epidemiol 44, 512–525, doi:dyv080 [pii] 10.1093/ije/dyv080 (2015).

82 Pierce, B. L. & Burgess, S. Efficient design for Mendelian randomization studies: subsample and 2-sample instrumental variable estimators. American journal of epidemiology 178, 1177–1184 (2013).

83 John, J., Kodama, T. & Siegel, J. M. Caffeine promotes glutamate and histamine release in the posterior hypothalamus. Am J Physiol Regul Integr Comp Physiol 307, R704–710, doi:ajpregu.00114.2014 (2014).

84 Wallace, C. H. R., Baczkó, I., Jones, L., Fercho, M. & Light, P. E. Inhibition of cardiac voltage-gated sodium channels by grape polyphenols. British journal of pharmacology 149, 657–665 (2006).

85 Wang, J.-H. et al. Modulation of Ca2+ signals by epigallocatechin-3-gallate (EGCG) in cultured rat hippocampal neurons. International journal of molecular sciences 12, 742–754 (2011).

